# Rictor induces AKT signaling to regulate lymphatic valve formation

**DOI:** 10.1101/2023.06.12.544698

**Authors:** Richa Banerjee, Luz A. Knauer, Drishya Iyer, Sara E. Barlow, Joshua P. Scallan, Ying Yang

## Abstract

Lymphatic valves are specialized structures of the collecting lymphatic vessels and are crucial for preventing retrograde lymph flow. Mutations in valve-forming genes have been clinically implicated in the pathology of congenital lymphedema. Lymphatic valves form when oscillatory shear stress (OSS) from lymph flow signals through the PI3K/AKT pathway to promote the transcription of valve-forming genes that trigger the growth and maintenance of lymphatic valves throughout life. Conventionally, in other tissue types, AKT activation requires dual kinase activity and the mammalian target of rapamycin complex 2 (mTORC2) commands this process by phosphorylating AKT at Ser473. Here we showed that embryonic and postnatal lymphatic deletion of *Rictor*, a critical component of mTORC2, led to a significant decrease in lymphatic valves and prevented the maturation of collecting lymphatic vessels. *RICTOR* knockdown in human lymphatic endothelial cells (hdLECs) not only significantly reduced the level of activated AKT and the expression of valve-forming genes under no-flow conditions, but also abolished the upregulation of AKT activity and valve-forming genes in response to flow. We further showed that the AKT target, FOXO1, a repressor of lymphatic valve formation, had increased nuclear activity in *Rictor* knockout mesenteric LECs, *in vivo*. Deletion of *Foxo1* in *Rictor* knockout mice restored the number of valves to control levels in both mesenteric and ear lymphatics. Our work revealed a novel role of RICTOR signaling in the mechanotransduction signaling pathway, wherein it activates AKT and prevents the nuclear accumulation of the valve repressor, FOXO1, which ultimately allows the formation and maintenance of a normal lymphatic valve.

## Introduction

The lymphatic vasculature is critical for maintaining tissue fluid homeostasis in the body. Lymphatic capillaries absorb interstitial fluids, immune cells, and lipids to form lymph and collecting lymphatic vessels transport lymph back to the bloodstream [1, 2]. The collecting lymphatic vessels ensure forward lymph flow by growing intraluminal bicuspid valves that prevent retrograde flow. The lymphatic valves are comprised of two lymphatic endothelial cell (LEC) layers separated by an extracellular matrix (ECM) core [3]. Defective lymphatic valves have been implicated in the human lymphatic disease, lymphedema, characterized by tissue swelling due to the accumulation of interstitial fluids, deposition of adipose tissue, fibrosis, and susceptibility to infections [4–6]. Mutations in many valve genes that are involved in human congenital lymphedema such as *FOXC2*, *GATA2, ITGA9*, *CX34* etc., cause valve defects in their associated mouse models [7–15].

The formation of lymphatic valves is regulated by the mechanotransduction signaling pathway. LECs at vessel branch points experience oscillatory shear stress (OSS) due to turbulent lymph flow and they respond by upregulating valve transcription factors like, Prox1, Foxc2, and Gata2 to form lymphatic valves [16]. OSS signals through VE-cadherin on the cell membrane of LECs, which activates the β-catenin and AKT cellular pathways to regulate valve formation and maintenance [17, 18]. As a key component in the mechanotransduction signaling pathway, the activity of AKT regulates many cellular processes, including lymphangiogenesis and lymphatic valve formation [19]. In support of this, injecting the AKT activating drug, SC-79, in WT mice was shown to induce ectopic lymphatic valve growth [17]. In many cell types, the activation of AKT requires the phosphorylation of two critical residues: threonine 308, which is phosphorylated by the protein dependent kinase 1 (PDK1) and serine 473, which is phosphorylated by the mammalian target of rapamycin complex 2 (mTORC2) [20]. mTORC2 is involved in many downstream cellular events such as cell migration, proliferation, survival, and cytoskeletal re-arrangement [20]. RICTOR is a critical component of the multi-protein mTORC2, which is required for the phosphorylation of Ser473 to fully activate AKT [21]. However, whether this function of RICTOR in AKT activation is important in the lymphatic vasculature remains unknown. Here we unveiled the role of *Rictor* in the formation of lymphatic valves. We found that lymphatic specific deletion of *Rictor* led to a significant loss of lymphatic valves in the mesenteric and ear lymphatic vessels. We provided evidence that *RICTOR* knockdown (KD) in cultured human dermal lymphatic endothelial cells (hdLECs) decreased AKT activation and subsequently the expression of important lymphatic valve genes. We revealed that *Rictor* knockout (KO) LECS, *in vivo*, expressed increased levels of nuclear FOXO1, a repressor of lymphatic valve formation which is a downstream target of AKT. Furthermore, we showed that ablation of *Foxo1* restored the number of valves to control levels in *Rictor* KO mice. Therefore, RICTOR induced AKT activation in the mechanotransduction signaling pathway is required for lymphatic valve formation.

## Results

### Embryonic deletion of Rictor leads to loss of valves in mesenteric lymphatic vessels

To investigate the unknown role of *Rictor*, a critical component of mTORC2, in lymphatic valve formation, we used a tamoxifen (TM) inducible LEC specific Cre line *Prox1CreER^T2^* [22], to conditionally delete exon 11 of *Rictor* (*Rictor^flox/flox^*) in embryonic mesenteric lymphatics, but not in the blood vasculature. To aid in visualizing the lymphatic vasculature, a LEC reporter line, *Prox1-GFP* [23] was crossed with the *Prox1CreER^T2^;Rictor^flox/flox^*mice to generate the *Prox1CreER^T2^;Rictor^flox/flox^;Prox1-GFP* mice (also referred to as *Rictor^LEC-KO^*). The *Rictor^flox/flox^;Prox1-GFP* littermates were administered the same dosage of TM and used as controls. Pregnant dams were injected with TM on embryonic day (E) 13.5 and E14.5 when the embryonic mesenteric lymphatic vessels begin to develop (Figure 1A). The mesenteries were analyzed on E18.5 when the lymphatic vessel network is established as well as the first timepoint when mature valves are observed (Figure 1A). Although there were no structural differences between the lymphatic vessels of *Rictor^LEC-KO^*and control embryos, *Rictor^LEC-KO^* mesenteric vessels had fewer GFP^high^ positive valves compared to the controls (Figure 1B and C, arrowheads). To confirm the reduction in GFP^high^ valves in *Rictor^LEC-KO^*mesentery, we measured the length (millimeter, mm) of each vessel and counted the number of GFP^high^ valves per vessel to generate valves per mm. *Rictor^LEC-KO^* mesenteries had 40% fewer valves per mm compared to control animals (*P*<0.05, Figure 1D). Whole-mount immunostaining for Prox1 revealed fewer Prox1^high^ valves in *Rictor^LEC-KO^*lymphatic vessels compared to control (Figure 1E and H, arrowheads). Valve LECs deposit fibronectin-EIIIA (FN-EIIIA) as part of their extracellular matrix core. Fibronectin receptor protein, integrin α9, is highly expressed in mature lymphatic valve leaflets and its deficiency leads to morphologically abnormal and dysfunctional lymphatic valves [8, 24, 25]. Whole-mount immunostaining revealed that although the *Rictor^LEC-KO^* mesentery possesses fewer valves compared to the control, the expression of integrin α9 is retained in the remaining valves (Figure 1F, G, I, and J). Together, RICTOR is needed for the formation of adequate lymphatic valves during embryonic development of the lymphatic vasculature.

**Figure 1.**
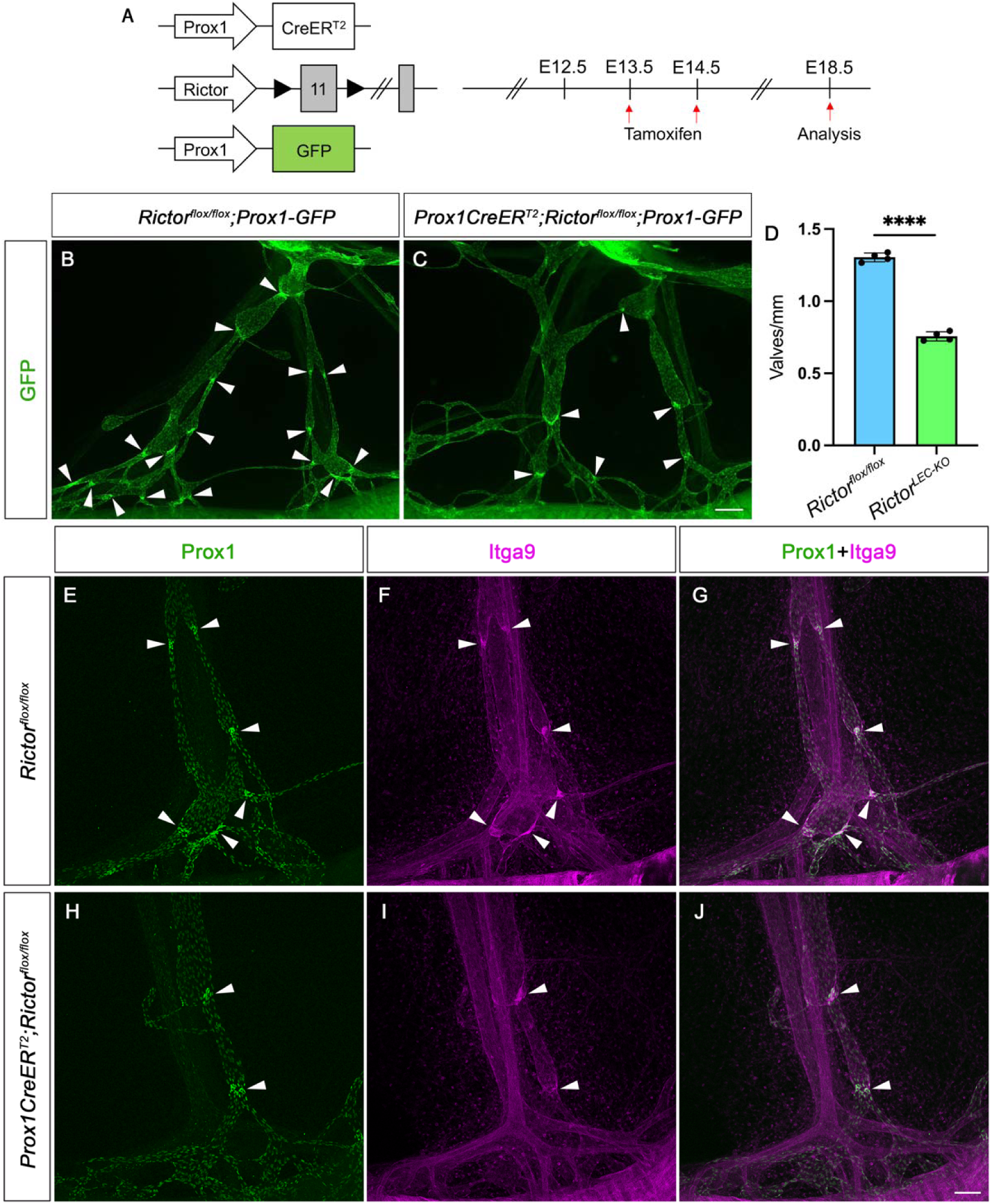
*Rictor* deletion leads to valve loss in embryonic mesentery. **A.** TM injection schedule for embryonic *Rictor* deletion. **B-C.** Fluorescence images of the mesenteric lymphatic vessels from the lymph node (top) to the gut (bottom) represented by GFP expression from E18.5 embryos. Arrowheads point to GFP^high^ valves. **D.** Valves per mm from each mesentery. **E-J.** Whole-mount immunostaining of E18.5 mesenteries post E13.5/E14.5 TM injections with Prox1 (green) and Itga9 (magenta). **G** is a merged image of **E** and **F**. **J** is a merged image of **H** and **I**. Arrowheads point to Prox1^high^ valves. Scale bars are 200 μm for B-C and 100 μm for E-J. All values are mean ± SEM. 4 control and 4 *Rictor^LEC-KO^* mesenteries were used for every analysis. \*\*\*\**P <* 0.0001, was calculated by unpaired Student’s *t* test.

### Ear lymphatic vessels lose valves and SMC coverage upon postnatal Rictor deletion

In addition to the embryonic deletion of *Rictor*, we induced lymphatic specific *Rictor* deletion at postnatal day (P) 1 and P3 (P1/P3 TM) via TM injection and analyzed the ear lymphatic vessels on P21 (Figure 2A). Since we were not able to find a working Rictor antibody to confirm deletion of the protein, we bred *Prox1CreER^T2^;Rosa26mT/mG* mice to verify the deletion efficiency and specificity our Cre line. To induce recombination, we injected TM at P1 and P3, the same TM injection schedule as in Figure 2A, and analyzed various tissues at P8, P14 and P21 (Supplemental Figure 1A). We found that Cre-induced GFP expression was present in all the lymphatic vessels in both mesentery and ears from various stages (Supplemental Figure 1B-BB). Immunostaining for Prox1 confirmed that the GFP signal was restricted to only Prox1-expressing LECs and absent in blood vessels (Supplemental Figure 1H-J). Since lymphatic valves in the ear begin developing after birth, it is an ideal tissue to investigate the role of Rictor in the formation of lymphatic valves postnatally. Like the embryonic mesentery, the *Rictor^LEC-KO^* ears showed fewer GFP^high^ valves in the collecting lymphatic vessels compared to the control ears (Figure 2B-E). Quantification revealed a 35% decrease in valves per mm in *Rictor^LEC-KO^* ears compared to control ears (*P*<0.05, Figure 2F). Since we deleted *Rictor* before the formation of lymphatic vessels and valves in the ear, our data revealed that *Rictor* regulates lymphatic valve formation, which is consistent with the results from embryonic deletion (Figure 1).

**Figure 2.**
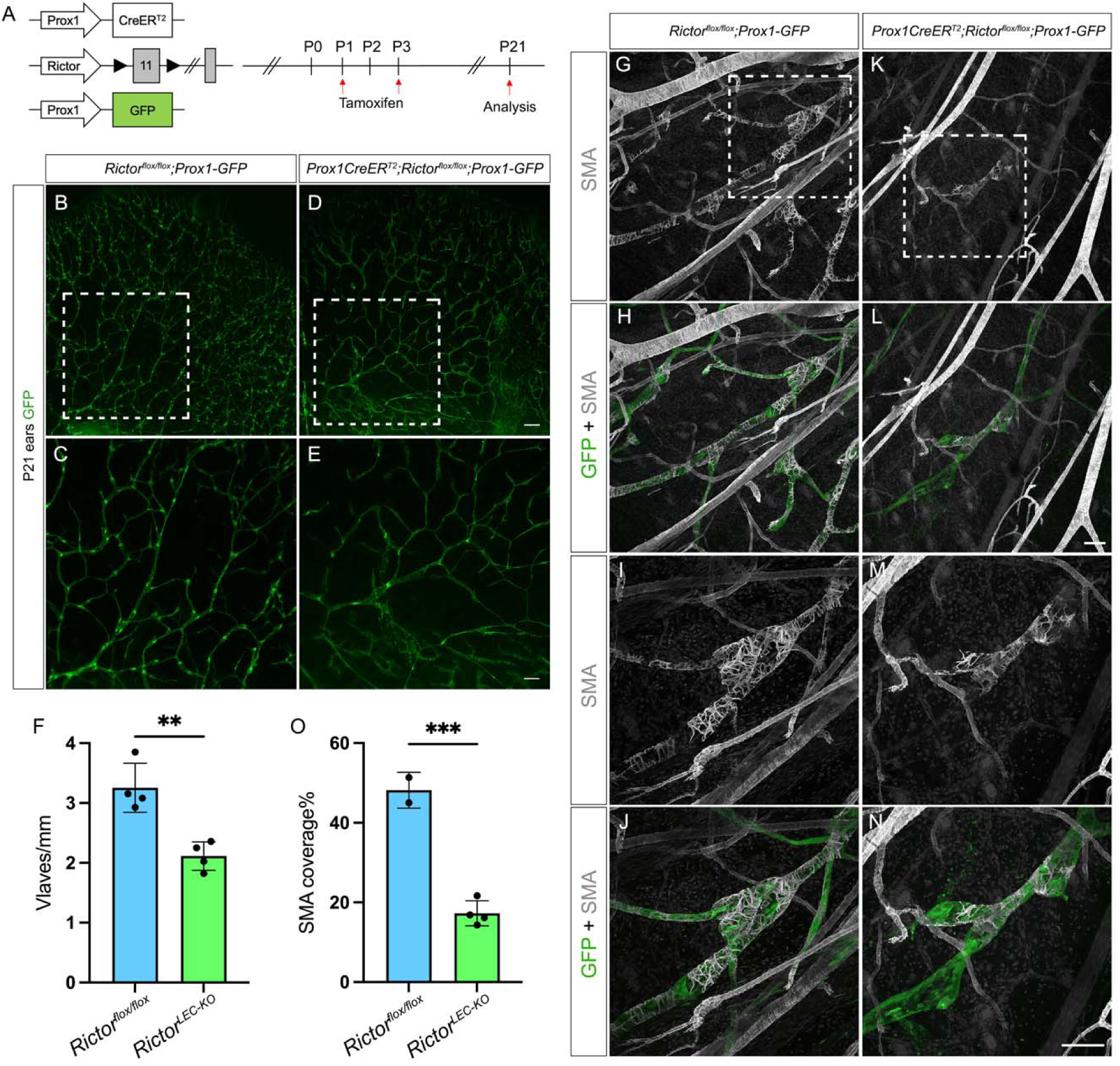
*Rictor* deletion leads to valve loss and reduced SMC coverage in the ear lymphatics. **A.** TM injection schedule for postnatal deletion of *Rictor*. **B-E.** Fluorescence images of the ear lymphatics from control (B and C) and *Rictor^LEC-KO^* (D and E) represented by GFP expression at P21. C and E are the higher magnification images of the white boxed region from B and D. **F.** Valves per mm from each mouse of control (blue bar) and *Rictor^LEC-KO^* (green bar). **G-N.** Whole-mount immunostaining of P21 ears with P1/P3 TM injection from control and *Rictor^LEC-KO^* with SMA (gray) and GFP (green). I, J, M and N are the higher magnification images of the white boxed region from G, H, K and L. **O.** Percentage of SMA coverage from each ear of control (blue bar) and *Rictor^LEC-KO^*(green bar). Scale bars are 500 μm for B and D, 200 μm for C and E, and 100 μm for G-N. All values are mean ± SEM. 4 control and 4 *Rictor^LEC-KO^* mice were used for every analysis. \*\**P <* 0.01 and \*\*\**P <* 0.001 were calculated by unpaired Student’s *t* test.

Since mTORC2 signaling is traditionally known to control cellular proliferation, we performed whole-mount immunostaining of P21 control and *Rictor^LEC-KO^* ear skin tissue with GFP and Ki67, a widely used proliferation marker [26] (Supplementary Figure 3A and B). Quantification of Ki67^+^/GFP^+^ double positive and all GFP^+^ positive LECs revealed no significant difference in the percentage of Ki67^+^/GFP^+^ double positive LECs between *Rictor* knockout and control animals (Supplementary Figure 3C). Hence, the loss of valves in *Rictor^LEC-KO^* tissues cannot be due to reduced LEC proliferation.

Since we found that Rictor regulates lymphatic valve development, we next wanted to investigate whether Rictor regulates other features of collecting lymphatic vessels. One of the hallmarks of collecting lymphatic vessel maturation is the recruitment of smooth muscle cells (SMCs) to the vessel wall, specifically to the lymphangion regions that are devoid of valves. [27]. To test whether Rictor regulates SMC recruitment, we performed whole-mount immunostaining for smooth muscle α-actin (SMA) in the P21 ear skin of *Rictor^LEC-KO^* and control animals, and subsequently quantified SMA coverage in the collecting lymphatic vessels. While the collecting lymphatic vessels in the control ear had abundant and continuous SMC coverage as revealed by SMA staining (Figure 2G-J), the *Rictor^LEC-KO^*collecting lymphatic vessels had sparse coverage of SMCs (Figure 2K-N). Quantification of the amount of SMA positive areas revealed a significant reduction in SMCs coverage on the *Rictor^LEC-KO^*ear collecting lymphatic vessels compared to the control (Figure 2O, 17% vs. 48%). This suggests that *Rictor* plays a role in SMC recruitment to the collecting lymphatic vessels during development and its deletion leads to incomplete vessel maturation.

To further investigate the defect in SMC recruitment post *Rictor* deletion, we examined the expression of different vascular SMC recruitment factors in cultured human dermal lymphatic endothelial cells (hdLECs) treated with scramble sh*RNA* (Scramble control) or sh*RNA* against *RICTOR* (sh*RICTOR*) and exposed them to flow (OSS) or no flow (static) conditions. We performed quantitative realtime PCR (qRT-PCR) to assess the expression of selected endothelial cell-derived SMC recruitment factors in control and *RICTOR* knockdown hdLECs. Although platelet-derived growth factor B (*PDGFB*) null mice have impaired SMC recruitment to lymphatic vessels [28], our qRT-PCR results showed *RICTOR* knockdown in hdLECs does not affect the expression of *PDGFB* (Supplemental Figure 2). We next analyzed the expression of several blood endothelial cell (BEC)-derived factors implicated in blood vascular pericyte recruitment including Platelet-derived growth factor D (*PDGFD*), endothelin-1 (*EDN1*), transforming growth factor beta-1 (*TGFβ1),* Angiopoietin-1 (*ANGPT-1*), and heparin binding-EGF like growth factor (*HB-EGF*) [29]. In addition, we investigated the expression of endothelial expressed sphingosine-1 phosphate receptor 1 (*S1PR1*), also called endothelial differentiation gene 1 (*EDG1*), because *Edg1^-/-^* mice exhibit incomplete vascular maturation and a defect in vascular SMC migration [30]. We also analyzed the expression of the SMAD family member 5, *SMAD5*, because homozygous *SMAD5-*null embryos displayed abnormal blood vascular development characterized by dilated blood vessels and reduced vascular SMC coverage [31]. The qRT-PCR analysis showed that 1) *PDGFD, TGFB1, ANGPT1,* and *S1PR1* are significantly downregulated in hdLECs treated with sh*RICTOR* without OSS, 2) *HB-EGF* is significantly upregulated upon sh*RICTOR* treatment with and without OSS, 3) *PDGFB*, *EDN1*, and *SMAD5* are not affected by *RICTOR* knockdown, and 4) Exposure to OSS downregulates the expression of *PDGFD*, *TGFB1*, and *ANGPT1* in control and sh*RICTOR* treated hdLECs (Supplemental Figure 2). Therefore, our data suggests that the expression of *PDGFD*, *TGB1*, *ANGPT1,* and *S1PR1* genes in LECs is regulated by mTORC2 signaling and may have potential roles in lymphatic SMC recruitment.

Since we observed defects in valve formation and in SMC recruitment to *Rictor^LEC-KO^* collecting lymphatic vessels, it suggested an inadequacy in the maturation of the collecting lymphatic vessels. During maturation, collecting vessel LECs downregulate the expression of Lyve1, the lymphatic vessel endothelial hyaluronan receptor, whereas capillary LECs retain high Lyve1 expression [2]. To further confirm that *Rictor^LEC-KO^* mice experience a defect in collecting lymphatic maturation, we performed whole-mount immunostaining to investigate the expression of Lyve1 in *Rictor^LEC-KO^* and control mice (Supplemental Figure 3D-K). The collecting lymphatic vessels of the P21 control ear skin, located in the basal area of the ear skin tissue, did not display any Lyve1 staining (Supplemental Figure 3H and I). In contrast, the intermediate area, which is the junction between the collecting lymphatic vessels and the capillaries, containing pre-collectors, displayed a gradual increase in Lyve1 expression in the control (Supplemental Figure 3D and E). In comparison to the control tissues, the collecting lymphatic vessels from the *Rictor^LEC-KO^* ear skin tissues began expressing Lyve1 at the basal area (Supplemental Figure 3J and K) and continued to express Lyve1 at the junction between the collecting lymphatic vessels and the capillaries (Supplemental Figure 3F and G). We found that 57% of collecting vessel LECs were Lyve1^+^ in *Rictor^LEC-KO^*mice and in contrast, only 8% of collecting vessel LECs were Lyve1^+^ in control animals (*P*<0.05, Supplemental Figure 3L). The data demonstrated that the collecting lymphatic vessels had a more capillary-like phenotype upon *Rictor* deletion and confirms that mTORC2 signaling is required for collecting lymphatic vessel maturation.

### Postnatal deletion of Rictor leads to valve loss in mesenteric lymphatic vessels

After lymphatic valves are formed, they require constant mechanotransduction signaling to maintain their structure throughout life [17]. To investigate the role of mTORC2 signaling in lymphatic valve formation and maintenance postnatally in mice, we induced the deletion of *Rictor* by administrating TM at P1 and P3 after the mesenteric lymphatic vessel network and valves are formed and the mesenteries were analyzed at P8, P14, and P21 (Figure 3A and Supplemental Figure 4A). The 100% recombination efficiency of the *Prox1CreER^T2^* was confirmed in *Prox1CreER^T2^;Rosa26^mT/mG^* mesenteric lymphatic vessels at P8, P14, and P21 (Supplemental Figure 1B-V). Although the overall morphology of the lymphatic vasculature was not altered upon *Rictor* deletion, the number of GFP^high^ valves were decreased in the *Rictor^LEC-KO^* mesentery compared to their littermate controls at all the three stages (Figure 3B-E, 3H-K and Supplemental Figure 4B-E, arrowheads). To confirm that valve loss is not due to the reduction in vessel length, we measured the length of each collecting vessel and its associated pre-collectors and normalized valve number to the vessel length (mm) to obtain the valves per mm. While the number of valves per mm was significantly lower in the *Rictor^LEC-KO^* mesentery compared to the control at P8, P14, and P21 (Figure 3F, 35%; 3L, 36%; and Supplemental Figure 4F, 35% decrease, *P*<0.05), the average length of each vessel (mm) was not significantly different between control and *Rictor^LEC-KO^*(Figure 3G, 3M, and Supplemental Figure 4G). Thus, in addition to embryonic valve development, RICTOR also regulates postnatal lymphatic valve formation and maintenance after birth.

**Figure 3.**
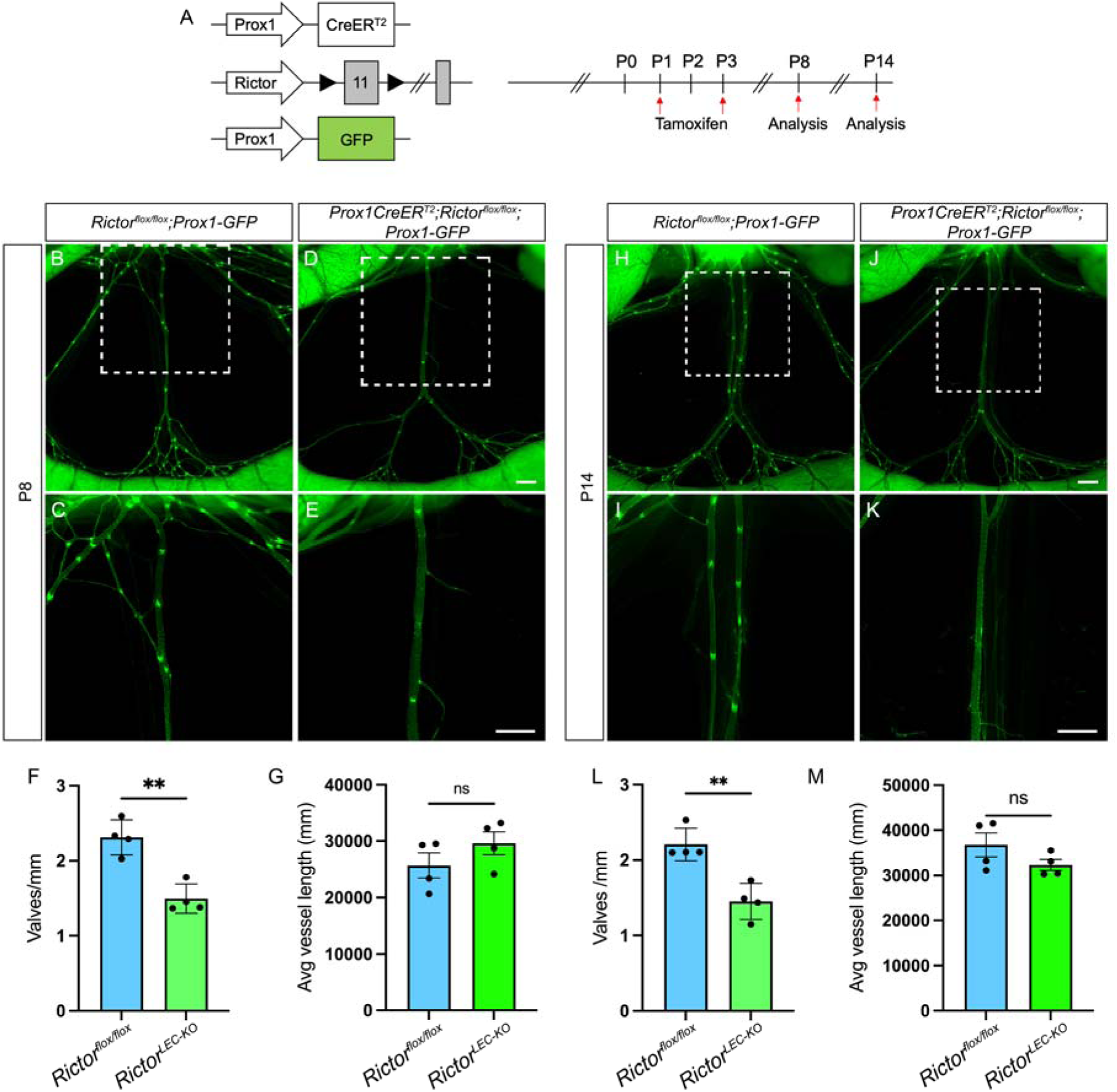
Postnatal deletion of *Rictor* leads to valve loss in the mesentery. **A.** TM injection procedure for postnatal deletion of *Rictor.* **B-E, H-K.** Fluorescence images of the mesenteric lymphatic vasculature represented by GFP expression from P8 (B-E) and P14 (H-K). C, E, I, and K are higher magnification images of the white squared areas from B, D, H, and J. **F and L**. Valves per mm from each mesentery of control (blue bar) and *Rictor^LEC-KO^* (green bar). **G and M**. Average vessel length of each vessel from control (blue bar) and *Rictor^LEC-KO^* (green bar). Scale bars are 500 μm in B-E and H-K. All values are mean ± SEM. 4 control and 4 *Rictor^LEC-KO^*mesenteries were used for the analysis. \*\**P <* 0.01 and ns=not significant, was calculated by unpaired Student’s *t* test.

### Valves in Rictor^LEC-KO^ mesentery retain the expression of valve genes and junctional proteins

To investigate whether the remaining valves in *Rictor^LEC-KO^*lymphatics retained the characteristics of wildtype valves, we performed whole-mount immunostaining on the P8 mesenteries with the same TM protocol as in Figure 3A to induce *Rictor* deletion. The expression of Prox1, a marker for lymphatic identity, remained unchanged in control and *Rictor^LEC-KO^* valves. However, the number of Prox1^high^ valves was evidently reduced in *Rictor^LEC-KO^* vessels (Figure 4A-D). Integrin α9, is highly expressed in mature valve LECs designating the complete bi-leaflet structure of a lymphatic valve. *Itga9* deficient mice develop chylothorax and die perinatally and this phenomenon is attributed specifically to morphologically abnormal and dysfunctional lymphatic valves [8, 24]. Like the control valves, *Rictor^LEC-KO^* valves expressed a high level of Itga9 (Figure 4E-H). The valve LECs express elevated level of Foxc2 compared to the lymphangion (vessel between two valves) LECs and Foxc2 expression induced by OSS is essential for lymphatic valve morphogenesis and maintenance [32, 33]. The valve LECs in *Rictor^LEC-KO^*still expressed a higher level of Foxc2 compared to the lymphangion LECs (Figure 4I-L), like the control. Junctional proteins are imperative in maintaining vessel integrity as well as serving as signaling molecules that drive downstream regulation of valve genes. VE-cadherin is an adherens junction protein, and it regulates valve development and integrity in response to OSS [17]. VE-cadherin (Cdh5) expression and localization appeared similar in LECs of *Rictor^LEC-KO^* and control (Figure 4M-P). Claudin 5 (Cldn5) is a tight junction protein and its expression is required for collecting lymphatic vessel integrity since *Cldn5^+/-^* mice have dilated and leaky lymphatics post an ultraviolet B irradiation challenge [34]. *Rictor^LEC-KO^* and control LECs showed no disparity in Cldn5 expression (Figure 4Q-T). In contrast to the ears, control and *Rictor^LEC-KO^* mesenteries displayed similar SMC wrapping patterns around the lymphangion (Figure 4U-X). In summary, these data showed that *Rictor^LEC-KO^* mesenteries maintained structurally intact lymphatic vessels with fewer valves.

**Figure 4.**
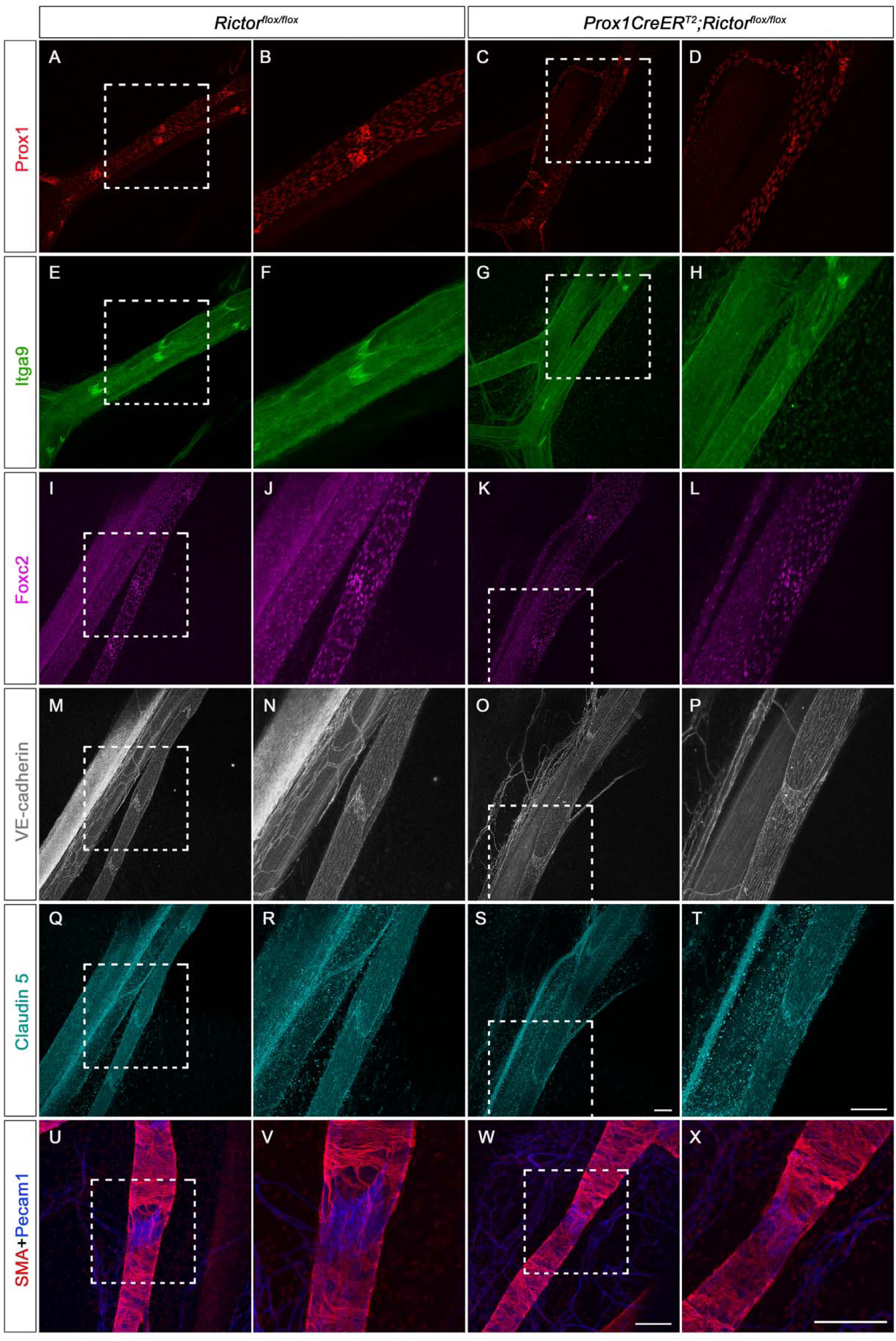
The expression of valve genes, junctional proteins, and SMA is present in the lymphatic vessels of *Rictor^LEC-KO^* mesentery with postnatal deletion. **A-X.** Whole-mount immunostaining of P8 mesenteries with P1/P3 TM injection from control and *Rictor^LEC-KO^*with Prox1 (red, A-D), Itga9 (green, E-H), Foxc2 (magenta, I-L), VE-cadherin (gray, M-P), Claudin 5 (cyan, Q-T), SMA (red, U-X), and Pecam1 (blue, U-X). B, D, F, H, J, L, N, P, R, T, V, and X are the higher magnification images of the white squared areas from A, C, E, G, I, K, M, O, Q, S, U, and W. Scale bars are 100 μm for A-X. 4 controls and 4 knockouts were used in the analysis.

### RICTOR knockdown eliminates the upregulation of valve genes in response to flow due to reduced AKT activity

Oscillatory lymph flow, which generates OSS, in response to which LECs signal via the mechano-transduction pathway to activate the expression of valve genes, such as *Prox1*, *Foxc2*, and *Gata2*, which leads to the formation of valves [16]. Since lymphatic-specific deletion of *Rictor* led to defective lymphatic valve formation and maintenance, we investigated the role of *Rictor* in regulating the expression of valve genes *in vitro*. To identify the transcriptional effects of *Rictor,* we treated cultured hdLECs with a control lentiviral sh*RNA* or a sh*RNA* to knockdown *RICTOR* and exposed the hdLECs to static/OSS condition post infection. We performed qRT-PCR to identify the expression of common valve and flow responsive genes like *PROX1*, *FOXC2, ITGA9*, *GJA4*, and *KLF4* in the four different groups: 1) hdLECs treated with Scramble sh*RNA* under static condition), 2) hdLECs treated with sh*RICTOR* under static condition, 3) hdLECs treated with Scramble sh*RNA* and OSS, and 4) hdLECs treated with sh*RICTOR* and OSS. We observed that the expression levels of *FOXC2, KLF4, ITGA9, PROX1,* and *NOS3* are significantly downregulated and the expression of *GJA4* (*connexin 37*) was almost abolished in the sh*RICTOR* treated cells under the static condition (Figure 5A). The increased expression of valve and flow responsive genes in the presence of OSS has been reported before [32]. Intriguingly, even in OSS, the expression levels of *FOXC2, KLF4, ITGA9, GJA4*, *PROX1* and, *NOS3* were significantly reduced in sh*RICTOR* treated hdLECs (Figure 5A). These valve and flow responsive genes showed a similar trend of expression at the protein level as well (Figure 5B). This shows that RICTOR is required for the upregulation of the critical valve and flow responsive genes in response to OSS. Since *Rictor* activates AKT through serine 473 phosphorylation in many cell types, we next investigated the activity of AKT upon *RICTOR* KD in LECs. Our western blot data showed that the level of Phospho-AKT (Ser473), a downstream target of mTORC2 signaling, was upregulated in response to 30’ OSS treatment (Figure 5C). However, this upregulation was eliminated in sh*RICTOR* treated hdLECs, which confirms that RICTOR regulates AKT activation in response to OSS in LECs (Figure 5C). In contrast, the levels of Phospho-AKT (Thr308) were not changed in response to sh*RICTOR* with or without flow treatment, which supports previous reports that PDK1, not RICTOR, activates AKT through threonine 308 phosphorylation.

**Figure 5.**
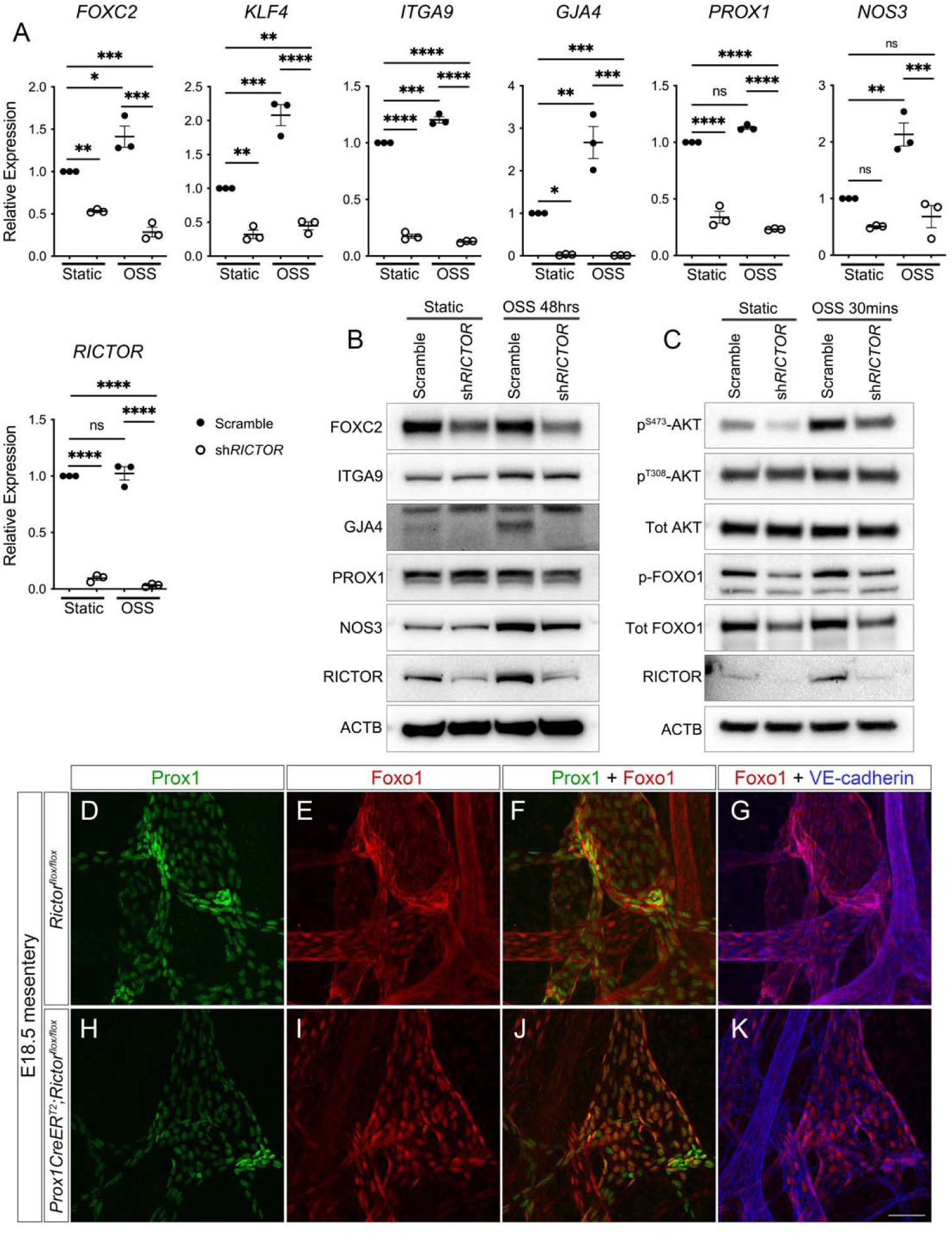
Loss of *Rictor* leads to downregulation in valve genes and increased nuclear FOXO1 due to reduced AKT activity. **A.** qRT-PCR performed for indicated genes in hdLECs transfected with a control scramble (closed circles) or sh*RICTOR* (open circles) and exposed to either static or OSS conditions (48 hours). **B.** Western blot was performed for the indicated genes using hdLECs transfected with a control scramble or sh*RICTOR* and exposed to either static or OSS conditions (48 hours). **C.** Western blot was performed in hdLECs transfected with a control scramble or sh*RICTOR* and exposed OSS conditions for 30 mins. **D.** Whole-mount immunostaining performed for PROX1 (green), FOXO1 (red), and VE-cadherin (blue) on E18.5 mesenteries (TM E13.5/14.5). Scale bars are 50 μm in D-K. 4 control and 4 *Rictor^LEC-KO^* embryonic mesenteries were used for the analysis in D-K**. \****P < 0.05, **P < 0.01, ***P < 0.001, ****P < 0.0001* and ns= not significant were calculated using two-way ANOVA with Tukey’s multiple comparisons in A.

### Rictor deletion enables Foxo1 localization in the LEC nucleus

Forkhead box protein O1 (FOXO1) is a member of the forkhead box O class (FoxO) subfamily of the forkhead transcription factors and is involved in glucose metabolism, cell proliferation, and tumor suppression [35]. Previously, we have shown that nuclear Foxo1 represses lymphatic valve genes and deletion of *Foxo1* embryonically and postnatally induces the formation of new valves [32]. AKT-induced phosphorylation of Foxo1 results in the nuclear export of Foxo1. This attenuates the Foxo1-mediated repression of valve genes and promotes valve gene expression [32]. Since we showed that AKT activity was downregulated in the loss of *Rictor*, we hypothesized that loss of *Rictor* results in increased nuclear retention of Foxo1. To test this, we performed whole-mount immunostaining of FOXO1 in E18.5 *Rictor^LEC-KO^*and littermate control mesenteries after inducing *Rictor* deletion in a pregnant dam by TM injections on E13.5 and E14.5 (Figure 5D-K). In control mesenteric LECs, Foxo1 staining is mostly in the cytoplasm (Figure 5E-G), however *Rictor^LEC-KO^* mesenteries showed more nuclear localization of Foxo1 staining (Figure 5I-K). Furthermore, western blot results from hdLECs treated with a scramble sh*RNA* or a sh*RNA* against *Rictor* (sh*RICTOR)* and exposed to flow or no flow conditions showed that the level of phosphorylated FOXO1 (p-FOXO1), which is non-nuclear FOXO1, was increased in response to flow. However, upon treatment with sh*RICTOR,* the level of p-FOXO1 protein declined (Figure 5C). These data confirm that RICTOR regulates cellular localization of Foxo1 through AKT activity.

Our data revealed that *Rictor* deletion leads to the increased localization of Foxo1 in the LEC nucleus and inhibits valve formation. To further support this finding, we generated a new mouse model with excessive Foxo1 in the nucleus. In this model, a mouse *Foxo1* cDNA in which the three main AKT phosphorylation sites, Threonine 32 (T32), Serine 253 (S253), and Serine 315 (S315), were changed to Alanine (A) by point mutations and this *Foxo1* cDNA with AAA mutations was knocked into the Rosa26 locus (Supplemental Figure 5A). A stop sequence was floxed by two Loxp sites to enable inducible expression of the *Foxo1*AAA cDNA by Cre recombinase. Thus, the overexpressed Foxo1 protein cannot be phosphorylated by AKT and excluded out of the nucleus. Therefore, this new mouse model harbors high nuclear activity of Foxo1. Additionally, an IRES sequence followed by tdTomato reporter was linked to the mutated *Foxo1*AAA cDNA, which resulted in a red fluorescence signal in the cells with overexpression of mutated Foxo1 protein. Therefore, this new mouse model is referred to as *Rosa26-Foxo1AAA-tdTomato* (Supplemental Figure 5A). We bred *Prox1CreER^T2^* with *Rosa26-Foxo1AAA-tdTomato* and *Prox1-GFP* reporter strain to generate the *Prox1CreER^T2^*;*Rosa26-Foxo1AAA-tdTomato*;*Prox1-GFP* mice for conditional overexpression of Foxo1AAA protein in LECs (henceforth referred to as *R26^LEC-Foxo1AAA^*). The litter mates without the *Prox1CreER^T2^* allele were used as control. Tamoxifen was injected at P1/P3 and mesentery tissues were analyzed at P6 (Supplemental Figure 5B and C). As we expected, the lymphatic vessels in the *R26^LEC-Foxo1AAA^* mesentery had fewer valves compared to the control (Supplemental Figure 5D-G, arrowheads). Quantification revealed a 65% decrease in valves per mm in the *R26^LEC-Foxo1AAA^* mice compared to control animals (*P*<0.05, Supplemental Figure 5H), which is consistent with a previous report using a different *Foxo1AAA* mouse model [36].These data strongly demonstrate that nuclear Foxo1 represses lymphatic valve formation, which explains the mechanism of how the accumulation of Foxo1 protein in the nucleus caused by *Rictor* deletion results in the loss of lymphatic valves.

### Homozygous deletion of Foxo1 in Rictor KO mouse model rescues the loss of valves in mesenteric lymphatics

We previously showed that LEC specific *Foxo1* ablation promotes the growth of new valves in wildtype mesenteric lymphatics and rescues valve loss in a mouse model of lymphedema [32]. Moreover, *Rictor* deletion increased the nuclear localization of Foxo1 in LECs *in vivo*, which led to significantly fewer valves embodied by the *R26^LEC-Foxo1AAA^* mice. Therefore, we hypothesized that deletion of *Foxo1* would promote the growth of new valves and rescue the valve loss observed in *Rictor^LEC-KO^* mice. We bred *Foxo1^flox/flox^*mice with *Rictor^flox/flox^* mice to create a double floxed mouse model (*Foxo1^flox/flox^;Rictor^flox/flox^*) and bred them with *Prox1CreER^T2^* mice to generate a double deletion mouse model. This method allowed us to delete one or two alleles of *Foxo1* to analyze the *Prox1CreER^T2^;Foxo1^+/flox^;Rictor^flox/flox^*and the *Prox1CreER^T2^;Foxo1^flox/flox^;Rictor^flox/flox^*mice. The same LEC reporter line, *Prox1-GFP* was combined with the *Prox1CreER^T2^;Foxo1^+/flox^;Rictor^flox/flox^*and *Prox1CreER^T2^;Foxo1^flox/flox^;Rictor^flox/flox^*mice to generate *Prox1CreER^T2^;Foxo1^+/flox^;Rictor^flox/flox^;Prox1-GFP* (*Foxo1^+/flox^;Rictor^LEC-KO^*) and *Prox1CreER^T2^;Foxo1 ^flox/flox^;Rictor^flox/flox^;Prox1-GFP* (*Foxo1;Rictor^LEC-KO^*) for visualization purposes. *Foxo1* and *Rictor* were deleted simultaneously in postnatal pups by P1/P3 TM injections (Figure 6A) and mesenteries were analyzed on P14 and P21 (Figure 6A-I and Supplemental Figure 6A-I). We normalized the valve number on each vessel to the length of each vessel and found that while the number of GFP^high^ valves per mm was significantly reduced in *Rictor^LEC-KO^* mesentery, double allele deletion of *Foxo1* restored the valves per mm to control levels (Figure 6D, E, H, I, and J and Supplemental Figure 6D, E, H, I, and J). Although one allele deletion of *Foxo1* did not significantly reverse the valve loss caused by *Rictor* deletion, valve number per mm was partially increased (Figure 6F, G, and J and Supplemental Figure F, G, and J). Moreover, double deletion of *Foxo1* and *Rictor* did not present any major defects in the mesenteric lymphatic vasculature in both *Foxo1^+/flox^;Rictor^LEC-KO^*and *Foxo1;Rictor^LEC-KO^* mice shown by no changes in average vessel length between the mice models (Figure 6K and Supplemental Figure 6K). To show that the new valves produced by the *Foxo1;Rictor^LEC-KO^* mesenteric lymphatic vessels were similar to the control valves, we performed whole-mount immunostaining of P14 control and *Foxo1;Rictor^LEC-KO^* mesenteries. Like the control mesenteric lymphatic valve LECs, the *Foxo1;Rictor^LEC-KO^* valves displayed brighter Foxc2, Prox1, and Itga9 staining compared to the lymphangion (Supplemental Figure 7A-P). Also, there was no distinction in Pecam1 and Claudin5 staining between the control and *Foxo1;Rictor^LEC-KO^* mesenteric lymphatic vessels (Supplemental Figure 7A-H). The *Foxo1;Rictor^LEC-KO^* mesenteric collecting lymphatic vessels also exhibited abundant SMC coverage on the lymphangion, like the control, represented by SMA staining (Supplemental Figure 7I-L). Thus, double-allele deletion of the valve repressor, *Foxo1*, was successful in promoting the growth of new and normal valves in *Rictor^LEC-KO^* mice mesentery.

**Figure 6.**
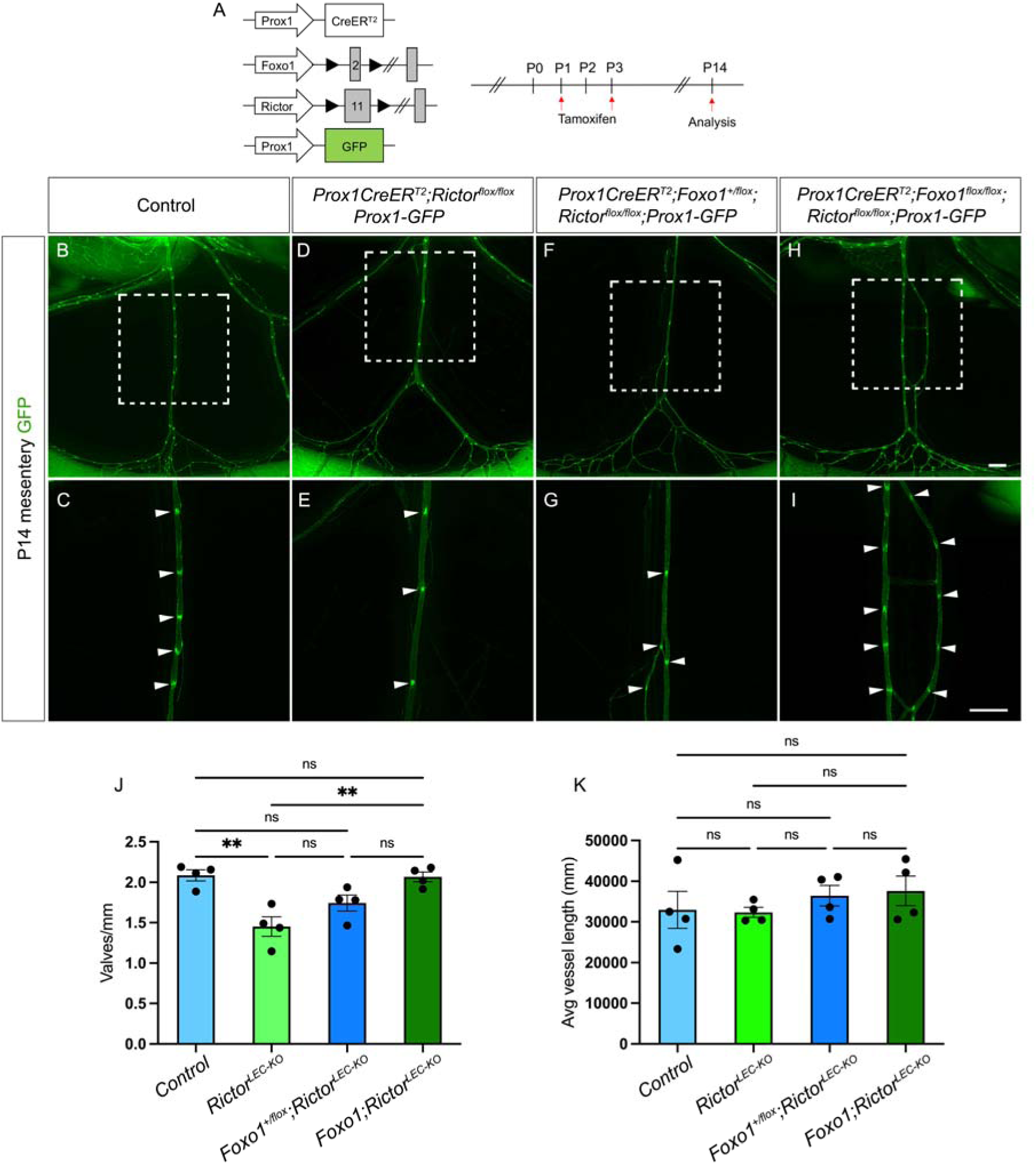
Postnatal deletion of *Foxo1* in mesentery rescues valve loss in *Rictor^LEC-KO^* mice. **A.** TM injection procedure for postnatal deletion of *Foxo1* and *Rictor.* **B-I.** Fluorescence images of the mesenteric lymphatic vasculature presented by GFP expression from P14. C, E, G, and I are higher magnification images of the white squared areas from B, D, F, and H. Arrowheads point towards GFP^high^ valves. **J**. Valves per mm from each mesentery: control (blue bar), *Rictor^LEC-KO^* (green bar), *Prox1CreER^T2^;Foxo1^+/flox^;Rictor^flox/flox^* (*Foxo1^+/flox^*;*Rictor^LEC-KO^*, dark blue bar), and *Prox1CreER^T2^;Foxo1 ^flox/flox^;Rictor^flox/flox^*(*Foxo1*;*Rictor^LEC-KO^*, dark green bar). Scale bars are 500 μm in B-I. All values are mean ± SEM. 4 controls and 4 knockouts were used in the analysis. \*\**P <* 0.01 and ns=not significant, was calculated by one-way ANOVA test.

### Postnatal single-allele deletion of Foxo1 restores valve loss in Rictor^LEC-KO^ mouse ear lymphatics

We have shown that deletion of *Foxo1* can restore valve loss in *Rictor^LEC-KO^* mesentery. Next, we tested our hypothesis that deletion of *Foxo1* in *Rictor^LEC-KO^*LECs ablates the excess Foxo1 that inhibits valve formation and protects against lymphatic valve loss in the ear tissue. *Rictor* and *Foxo1* were deleted by TM injections on P1 and P3 and the ears were analyzed on P21 (Figure 7A). In contrast to the results from the mesentery, one allele deletion of *Foxo1* in the *Foxo1^+/flox^;Rictor^LEC-KO^*ear lymphatics had more GFP^high^ valves compared to either the *Rictor^LEC-KO^* or two alleles deletion of *Foxo1* in the *Foxo1;Rictor^LEC-KO^*ears (Figure 7B-I). Additionally, the *Foxo1;Rictor^LEC-KO^* ear lymphatic vessels appeared more dilated than the vessels from the control, the *Rictor^LEC-KO^*, and the *Foxo1^+/flox^;Rictor^LEC-KO^* mice (Figure 7B-I). We quantified the valve number per mm and vessel diameter from these four genotypes. Our data showed that: 1) one allele deletion of *Foxo1* (*Foxo1^+/flox^;Rictor^LEC-KO^*) in the ears did not cause a significant increase in vessel diameter compared to the control (Figure 7K); 2) one allele deletion of *Foxo1* (*Foxo1^+/flox^;Rictor^LEC-KO^*) in the ears led to a significant increase in valves per mm compared to the *Rictor^LEC-KO^* ears and restored the valve number to control levels (Figure 7J); 3) two alleles deletion of *Foxo1* (*Foxo1;Rictor^LEC-KO^*) resulted in a grossly significant increase in vessel diameter along with significantly fewer valves per mm (Figure 7J and K) compared to the control and *Foxo1^+/flox^;Rictor^LEC-KO^* groups. The dilation of ear vessels in two alleles deletion of *Foxo1* was consistent with our previous data due to increased proliferation caused by *Foxo1* inactivation during development [32]. Additionally, we performed whole-mount immunostaining of SMA on control, *Rictor^LEC-KO^*, *Foxo1^+/flox^;Rictor^LEC-KO^* and *Foxo1;Rictor^LEC-KO^* ears and observed that SMC coverage is similar on control and *Foxo1^+/flox^;Rictor^LEC-KO^* lymphatic vessels (Figure 7L, M, P, and Q). However, SMA coverage is reduced on *Rictor^LEC-KO^* and *Foxo1;Rictor^LEC-KO^*lymphatic vessels (Figure 7N, O, R, and S). We quantified the SMA coverage on ear lymphatic vessels and showed that one allele deletion of *Foxo1* in *Rictor^LEC-KO^*ears restored the SMC coverage level to control levels (Figure 7T). However, two alleles deletion of *Foxo1* in *Rictor^LEC-KO^*ears did not rescue the loss of SMC coverage in *Rictor^LEC-KO^* ear lymphatic vessels (Figure 7T). This data demonstrates that *Foxo1* is a key repressor of lymphatic valves downstream of *Rictor* and a single allele deletion of *Foxo1* at a developmental stage is sufficient to rescue valve loss and SMC recruitment in the absence of *Rictor*.

**Figure 7.**
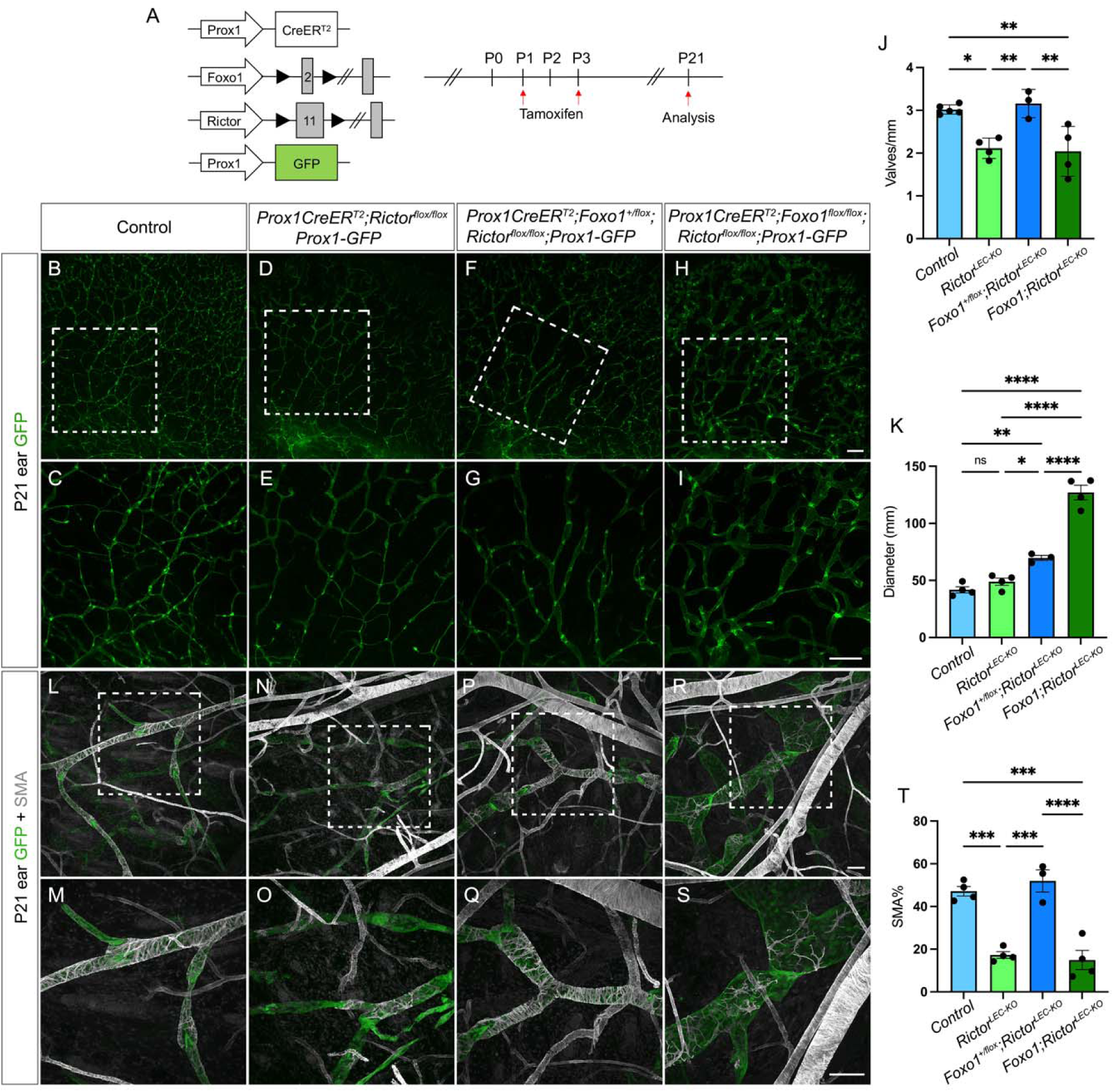
Postnatal *Foxo1* deletion rescues valve defect in *Rictor^LEC-KO^* ears. **A.** TM injection schedule to delete *Foxo1* and *Rictor*. **B-I**. Fluorescence images of P21 ear lymphatic vessels identified with GFP expression. C, E, G, and I are higher magnification images of white squared areas in B, D, F, and H. **J**. Valves per mm from control (blue bar), *Rictor^LEC-KO^*(green bar), *Prox1CreER^T2^;Foxo1^+/flox^;Rictor^flox/flox^*(*Foxo1^+/flox^*; *Rictor^LEC-KO^*, dark blue bar), and *Prox1CreER^T2^;Foxo1 ^flox/flox^;Rictor^flox/flox^* (*Foxo1*;*Rictor^LEC-KO^*, dark green bar). K. Diameter of collecting lymphatic vessels from each ear of control (blue bar), *Rictor^LEC-KO^* (green bar), *Prox1CreER^T2^;Foxo1^+/flox^;Rictor^flox/flox^*(*Foxo1^+/flox^*;*Rictor^LEC-KO^*, dark blue bar), and *Prox1CreER^T2^;Foxo1 ^flox/flox^;Rictor^flox/flox^*(*Foxo1*;*Rictor^LEC-KO^*, dark green bar). **L-S**. Whole mount-immunostaining of P21 ears (TM P1/P3) with SMA (grey) and GFP (green). M, O, Q, and S are higher magnification images of the white squared areas in L, N, P, and R. **T**. Percentage of SMA coverage from each ear of control (blue bar), *Rictor^LEC-KO^* (green bar), *Prox1CreER^T2^;Foxo1^+/flox^;Rictor^flox/flox^*(*Foxo1^+/flox^*;*Rictor^LEC-KO^*, dark blue bar), and *Prox1CreER^T2^;Foxo1 ^flox/flox^;Rictor^flox/flox^* (*Foxo1*;*Rictor^LEC-KO^*, dark green bar).. Scale bars are 500 μm for B-I and 100 μm for L-S. All values are mean ± SEM. 4 control and 4 KO mice were used for every analysis. \**P < 0.05,* \*\**P <* 0.01, \*\*\**P <* 0.001, \*\*\*\**P <* 0.0001, and ns=non-significant were calculated by one-way ANOVA tests in J, K, and T.

## Discussion

In our previous study, we showed that AKT activity was increased in response to OSS in LECs, which is critical for the upregulation of valve-forming genes in the mechanotransduction signaling pathway. Additionally, exogenous stimulation of AKT led to the production of new lymphatic valve s in wildtype mice [17]. These data are supported by a previous report that AKT is important for the formation of lymphatic valve s (Zhou et al. 2010). However, what is responsible for the activation of AKT in the mechanotransduction signaling pathway that stimulates lymphatic valve formation is thus far unknown. Although in other cell types the mTORC2 signaling is involved in the phosphorylation and activation of AKT [37], the role of mTORC2 signaling in the lymphatic vasculature, particularly in valve formation is yet untested. Here, we discovered a novel role of *Rictor*, a crucial component of mTORC2, in the regulation of lymphatic valve formation and maintenance. *Rictor* assembles the mTORC2 components and prosecutes downstream kinase activity. We showed that llymphatic specific deletion of *Rictor* led to a significant reduction in valve number embryonically and postnatally. Moreover, loss of *Rictor* severely affected the activity of AKT in OSS condition which, in turn, resulted in the decreased expression levels of valve-forming genes and increased nuclear activity of FOXO1, a downstream target of AKT. FOXO1 is a potent lymphatic valve repressor and by genetically ablating the expression of *Foxo1*, valve number was restored in *Rictor^LEC-KO^* mice. Therefore, our study revealed the role of the OSS-RICTOR-AKT-FOXO1 signaling axis in lymphatic valve formation (Figure 8).

**Figure 8.**
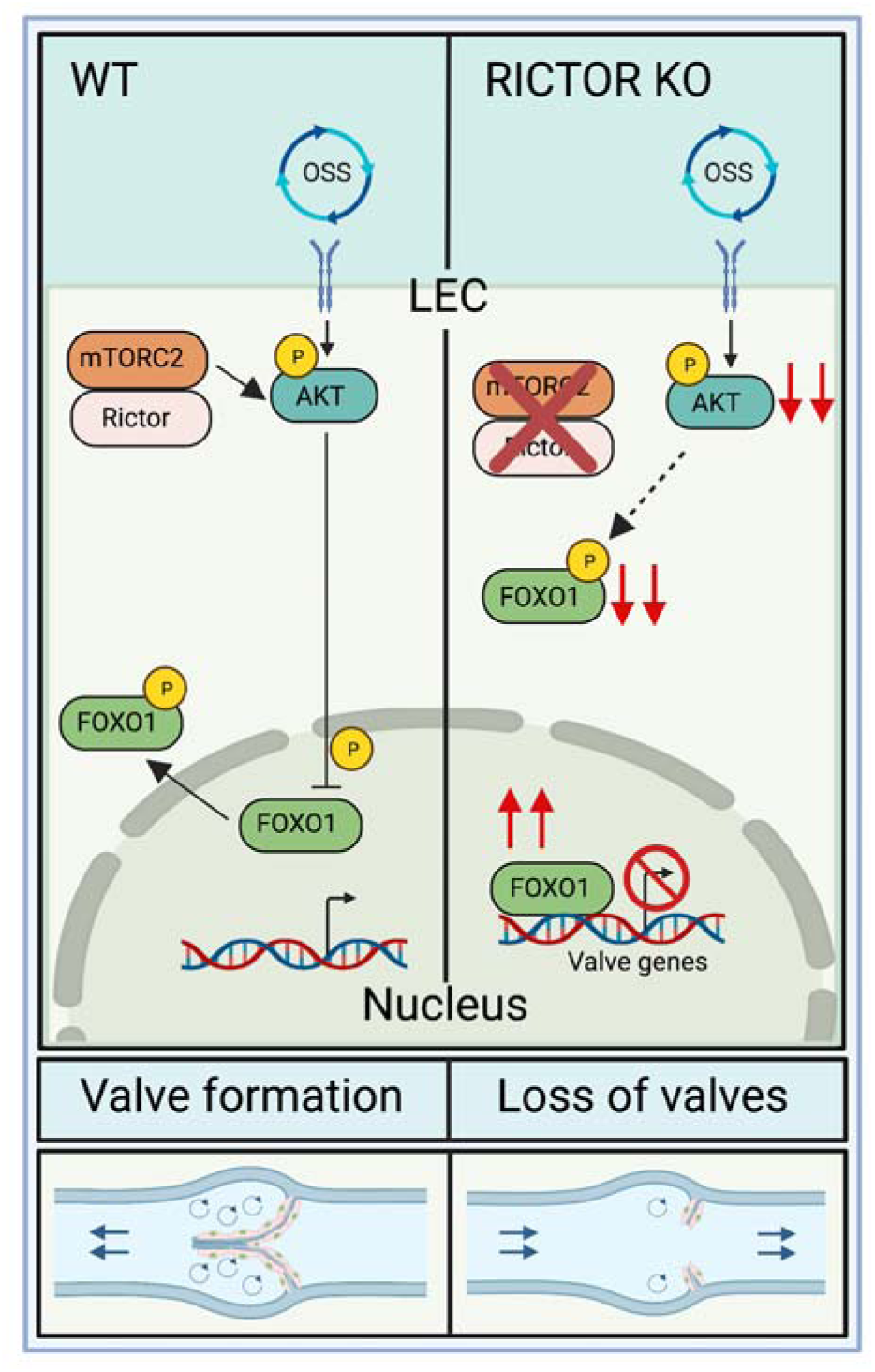
Rictor, a critical component of mTORC2, regulates lymphatic valve formation and maintenance via modulating AKT activation. In wildtype LECs, Rictor commands the complete activation of AKT which phosphorylates its downstream target, FOXO1, and expels it from the LEC nucleus, leading to the production of lymphatic valves. However, upon loss of Rictor signaling, AKT activation is inadequate to prevent FOXO1, which is a potent valve repressor, from executing its inhibitory function, thereby causing lymphatic valve loss.

### Rictor regulates lymphatic valve formation through AKT induced valve gene expression and nuclear Foxo1 inhibition

Lymph flow is oscillatory at branchpoints of the embryonic lymphatic vasculature and this oscillatory lymph flow stimulates a mechanotransduction signaling process that initiates the activation of AKT and leads to nuclear deport of the lymphatic valve repressor, FOXO1 [32]. This allows the transcriptional upregulation of important valve genes like *Foxc2*, *Gata2*, *Prox1*, *Itga9,* and *Gja4* (*Cx37*) and flow responsive genes like *Klf4* and *Klf2*, which causes the production and maintenance of lymphatic valves throughout life [2, 10, 38]. In this study, our data revealed that mTORC2 is upstream of AKT in the mechanotransduction signaling pathway and its critical component, *RICTOR*, is activated by oscillatory flow during lymphatic valve formation, which then phosphorylates AKT, at Ser473, to set forth the exit of nuclear FOXO1. We showed that when mTORC2 signaling was abolished by deletion of *Rictor,* there were fewer lymphatic valves in collecting lymphatic vessels *in vivo* and this loss of lymphatic valves was tissue type and developmental stage independent. Moreover, we revealed that the expression of important lymphatic valve genes and flow response genes (e.g FOXC2, ITGA9, GJA4, and KLF4) were curtailed in the absence of RICTOR in cultured hdLECs *in vitro*. We showed that the main cause of these phenotypes is due to the increased activity of Foxo1 in the LEC nucleus because nuclear Foxo1 inhibits the transcription of lymphatic valve genes, like Foxc2, by directly binding to its upstream promoter [32]. Therefore, valve formation was attenuated in our LEC specific Foxo1 nuclear overexpression mouse model, *R26^LEC-Foxo1AAA^*, which confirms that increased nuclear *Foxo1* inhibits the formation of lymphatic valves. These data prove that RICTOR induced AKT signaling is involved in the mechanotransduction pathway of lymphatic valve formation and maintenance. Interestingly, it is not surprising that Rictor is stimulated by OSS. Although in many cell types, the activity of mTORC2 signaling and AKT is increased in response to energetic stress, such as ligand dependent activation by IGF (insulin growth factor) stimulation, it has also been shown that mTORC2 signaling is stimulated by the tension built in focal adhesion points in mesenchymal stem cells (MSCs) due to dynamic mechanical signals experienced during tissue loading [39]. Our data showed that the expression levels of RICTOR in cultured hdLECs was increased with both 30 mins and 48 hrs OSS treatment, indicating that mTORC2 signaling is activated by physiological shear stress in the lymphatic vasculature in a ligand independent manner leading to the increase of activated AKT and lymphatic valve formation. However, there are still unanswered questions, for example, is RICTOR the only kinase in the mTORC2 complex that phosphorylates AKT? In our postnatal lymphatic *Rictor* knockout mouse model, the percentage of valve loss was maintained at ∼40% up to three weeks after deletion, which was very similar to the percentage of valve loss in *Foxc2* heterozygous mice. Although the *Rosa26^mT/mG^* reporter assay indicated 100% deletion of *Rictor*, valve number in *Rictor* LEC knockout mice never reduced to zero. Additionally, our whole-mount immunostaining data of P8 mesenteries exhibited that the remaining lymphatic valves in the *Rictor^LEC-KO^* lymphatic vasculature had the usual pattern of high expression of lymphatic valve genes (Foxc2, Prox1, and Itga9,) in the valves compared to the lymphangion. This suggests that although mTORC2 signaling is critical for lymphatic valve formation, there are other factors upstream or downstream of mTORC2 that influence lymphatic valve production and maintenance. Furthermore, despite that RICTOR was almost completely knocked down in cultured hdLECs, the level of phosphorylated AKT at Ser347 was not abolished but remained at a low level. Therefore, it is likely that other protein kinases in addition to RICTOR play a role in AKT activation in response to OSS.

### Rictor regulates collecting lymphatic vessel maturation

The mammalian lymphatic vasculature matures to form two distinct types of lymphatic vessels: lymphatic capillaries and collecting lymphatic vessels. Lymphatic capillaries express high levels of LYVE1 and collecting lymphatic vessels switch off the expression of LYVE1 during their maturation process [2]. Additionally, collecting lymphatic vessels recruit abundant SMCs to ensure the pulsatile contractile functioning whereas lymphatic capillaries are blind ended sacs that do no recruit SMCs [28]. We showed that upon postnatal *Rictor* deletion in mice, the SMC coverage on the collecting lymphatic vessels was severely reduced on the ear lymphatics but not on the mesenteric lymphatics. The mouse ear lymphatics develop after birth and our results indicate that ablating *Rictor* post birth negatively affects SMC recruitment to the collecting lymphatic vessels during their formation. Moreover, these collecting lymphatic vessels in the ear expressed abnormally high levels of Lyve1 while the mature collecting lymphatic vessels in control mice did not express Lyve1. Furthermore, we found that *RICTOR* knockdown in cultured hdLECs decreased the expression of many blood endothelial cell expressed SMC recruitment factors, including *PDGFD, TGFB1, ANGPT1, HBEGF,* and *S1PR1* [40–43]. Surprisingly, we observed no change in *PDGFB* expression *in vitro* upon *RICTOR* knockdown, although it has been reported that LEC specific *Pdgfb* knockout *in vivo* leads to a failure in SMC recruitment [28]. Similarly, *Edn1* plays a role in vascular SMC recruitment, however, its expression level did not change in *RICTOR* knockdown LEC either. Heparin-binding EGF-like growth factor (HB-EGF) is a ligand of the EGFR in endothelial cells and signals via the PI3K pathway to regulate vascular smooth muscle cell proliferation [44]. The unexpected upregulation in the expression of *HBEGF* upon *RICTOR* knockdown in cultured hdLECs can be explained as a feedback mechanism. HB-EGF binds to EGFR upstream of mTORC2 and its elevated expression upon *RICTOR* knockdown attempts to bring back the cells into homeostasis. Nevertheless, these data indicate that the maturation of collecting lymphatic vessels requires the activity of mTORC2 signaling. Ablation of *Rictor*, therefore, resulted in the excessive expression of Lyve1, the reduction of SMC recruitment, and decreased lymphatic valve formation in collecting lymphatic vessels.

### Postnatal deletion of Foxo1 rescues the valve loss in Rictor KO mice

Since the *R26^LEC-Foxo1AAA^* mouse model confirmed that excess nuclear Foxo1 prevents lymphatic valve formation, we hypothesized that *Foxo1* deletion in *Rictor^LEC-KO^* mice will abolish the repressive activity of Foxo1 and advocate the growth of valves to control levels. We showed that a single allele deletion of *Foxo1* induced new valves to grow in *Rictor^LEC-KO^* mice. Although valve number was not significantly increased in *Rictor^LEC-KO^* mice with one allele deletion of *Foxo1*, double allele deletion of *Foxo1* completely and significantly rescued the lymphatic valve loss in *Rictor* KO mesenteries. In contrast, the reduced number of valves seen in *Rictor* KO ear skin tissue only required a single allele deletion of *Foxo1* to be restored to the wildtype level. These data suggest that the effect of *Foxo1* inhibition works in a developmental stage dependent manner. In mice, mesentery develops embryonically and by the time the postnatal deletion of *Foxo1* and *Rictor* is induced, the lymphatic network in the mesentery has formed with fully developed lymphatic valves. On the other hand, since the ear lymphatics develop after birth, the postnatal deletion of *Foxo1* and *Rictor* occurs before these vessels form. Thus, our results suggest that LECs from embryos and from postnatal pups have different amounts of AKT activity. In other words, higher AKT activity is present in embryonic LECs compared to postnatal LECs. Therefore, more Foxo1 is needed to be out of the nucleus in postnatal vessels compared to embryonic ones to stimulate lymphatic valve formation. Hence, two allele deletion of *Foxo1* was required to reverse the valve loss observed in *Rictor^LEC-KO^*postnatal mesentery tissue while one allele deletion of *Foxo1* was enough to induce the growth of lymphatic valves in the ears. Furthermore, the lack of a proper amount of nuclear Foxo1 activity caused by the double allele deletion of *Foxo1* drove the dilation of the collecting lymphatic vessels in the ears, which led to a defect in SMC recruitment [32].

### AKT signaling yields many pathologically relevant clinical targets

Phosphoinositide 3-kinase (PI3K) mediated AKT (also called protein kinase B) signaling was thought to be critical for cell survival only but was later presented as a multifaceted protein kinase pathway involved in glycogen metabolism, cell cycle regulation and cellular senescence, and protein synthesis [45]. In blood endothelial cells (BECs), growth factors induced AKT signaling inhibits cell detachment induced apoptosis during angiogenesis and enhances migration by re-arranging actin cytoskeleton during angiogenic capillary formation [46–48]. In established blood vasculature, *Akt1* deletion in mouse endothelial cells causes a loss of vascular smooth muscle cell (VSMC) differentiation and leads to decreased arterial VSMC coverage, coronary structural deficiencies, and potent cardiac dysfunction [49]. Additionally, global *Akt1* knockout mouse models display defects in lymphangiogenesis, lymphatic valve formation, and SMC coverage on collecting lymphatics [19]. Many other mediators related to AKT signaling have been shown to cause lymphedema in murine mutation models. Protein tyrosine phosphatases (PTPs) work in conjunction with protein tyrosine kinases to modulate receptor signaling [50]. *PTPN11,* a type of intracellular PTP is downstream of many RTKs, including growth factor receptors and its mutation is associated with Noonan syndrome in humans [51]. Noonan syndrome is a complex disorder comprising facial dysmorphology, congenital heart disease and growth failure, and lymphedema is a clinical manifestation attributed to lymphatic dysplasia [52] and *PTPN11* mutations are also connected to chylothorax, lymphatic dilation, and lymph reflux in humans [53]. Interestingly PTPN11, also called SHP-2, negatively regulates the phosphorylation of VE-cadherin in HUVECs *in vitro,* and since VE-cadherin is upstream of AKT signaling in the lymphatic vasculature, it can be well hypothesized that *PTPN11* signals upstream of AKT in the lymphatic vasculature [54]. Another example is the protein tyrosine phosphatase non-receptor type-14 *(Ptpn14*) gene which encodes the Ptnpn14 enzyme in mice. Ptpn14 was shown to interact with Vegfr3 at the protein level and its gene trap mouse model displayed lymphatic hyperplasia and lymphedema in upper and lower limbs and periorbital region of adult mice [55]. Another upstream phosphatase of the PI3K-AKT signaling, Phosphatase and tensin homolog (PTEN), is known to reverse the phosphorylated and active phosphatidylinositol-3, 4, 5-triphosphate (PIP_3_) to inactive phosphatidylinositol-4, 5-bisphosphate (PIP_2_) and decrease AKT phosphorylation. When this inhibitory effect of PTEN is abolished in LEC specific *Pten* KO mice models, it results in increased lymphatic density in all major tissues and increased levels of phosphorylated *Akt* even though the level total *Akt* level is the same. LEC specific *Pten* KO mice also display reduced circulating inflammatory mediators and vessel leakiness. Although majority of the data is related to the role of PTPs in lymphnagiogenesis and very little is known about their direct effects on lymphatic valves, it is reasonable to postulate that these upstream signaling mediators of the AKT pathway are critical for the formation and maturation of lymphatic vessels which eventually experience oscillatory flow and develop lymphatic valves.

In summary, our finding is critical because it reveals a novel role of mTORC2 in the mechanotransduction signaling pathway that regulates lymphatic valve formation and maintenance. Further evidence for the significance of our findings is due to the clinical cases of cancer and organ transplant patients. These patients are administered mTOR inhibitors like, rapamycin and its orthologues, sirolimus and everolimus, and some of these patients develop lymphedema after long-time treatment [56–58]. Although the mammalian target of rapamycin complex 1 (mTORC1) is the direct target of rapamycin, prolonged exposure to rapamycin can affect the activity of mTORC2 [20]. Therefore, understanding the role of mTORC2-AKT signaling pathway in stimulating new valves to grow can help to develop therapy to treat valve defects in lymphedema.

## Materials and Methods

### Animals

Mouse husbandry and experiments were conducted under the guidelines of University of South Florida (USF) Institution Animal Care and Use Committee (IACUC). The *Rictor^flox/flox^*and *Foxo1^flox/flox^* mice were obtained from the Jackson laboratory. The use of *Foxo1^flox/flox^* and *Prox1CreER^T2^*mice have been described before [32, 59–62]. The *Prox1CreER^T2^*and *Prox1-GFP* mouse models were procured through the material transfer agreement (MTA). The *R26^LEC-Foxo1AAA^* mouse model was generated by Ingenious. All the mouse models were maintained a mixed genetic background (C57BL/6J x FVB) and both females and males were used. Embryonic deletion of *Rictor* was achieved by two intraperitoneal injections of TM (5 mg) into pregnant dams on E13.5 and E14.5. Postnatal deletion of *Rictor* and *Foxo1* was induced by injecting pups with 100 μg TM on P1 and P3.

### Ex vivo analysis of lymphatic vessel length, lymphatic valve, and vessel diameter quantification

Pregnant dams and *Prox1-GFP* reporter expressing postnatal pups were euthanized using IACUC approved methods and analyzed at the indicated experimental timepoint. To collect embryos, the uterine horn was detached from the dam and placed in cold phosphate buffered saline (PBS). Embryonic mesenteries were pinned *ex vivo* on a homemade Sylgard 184 pad. For postnatal pups, the mesentery was imaged *in situ* (without being detached from the body). Both embryonic and postnatal mesenteries were imaged under a Zeiss V16 microscope. The length of GFP positive large collecting lymphatic vessels appearing from the lymph node and the thinner branched pre-collector vessels towards the intestinal wall [32] (Figure 1B and C and Figure 3) were measured using the segmented line tool in Fiji (ImageJ) and the GFP^high^ valves were counted using the multi-point tool in Fiji. Ears from postnatal pups were excised from the mouse body at the indicated timepoint. The inner dermal layer was separated from the outer collagen layer. After the dermal layer was pinned on a sylgard 184 pad, it was analyzed under a Zeiss V16 microscope. In the ear, collecting lymphatic vessels towards the base of the ear were measured using the line tool in Fiji and GFP^high^ valves were counted using the multi-point tool in Fiji. The diameter of lymphatic vessels was counted using the line tool in Fiji. 4 KO mice and 4 littermate controls were used for each experiment.

### Quantification of cell proliferation

P21 ear skin tissue was stained with GFP and Ki67 and confocal images were acquired. GFP and Ki67 double positive LECs were counted using the multi-point tool using Fiji. The percentage of Ki67 positive cells were obtained by dividing the Ki67^+^ cells by the GFP^+^ cells multiplying the ratio by 100. 4 mice from each group were used for analysis.

### Smooth muscle cell coverage and LYVE1-expression area quantification

GFP/SMA or GFP/LYVE1 stained slides were imaged under a Zeiss V16 microscope at 32X and 25X magnification, respectively. The GFP/SMA or GFP/LYVE1 images were merged using the color and merge tool in Fiji (ImageJ). The areas of GFP positive collecting lymphatic vessels, avoiding the SMC rich blood vessels in GFP/SMA and LYVE1 positive capillaries in the GFP/LYVE1 tissues, were measured using the polygonal tool on Fiji and the areas were saved as a region of interest (ROI). The ROI was then applied to individual GFP and SMA or GFP and LYVE1 V16 images and the threshold of color was set using the image and adjust threshold option in Fiji. The threshold allows the measurement of area fraction of staining, which is then totaled individually. SMA% is obtained by dividing the total SMA area by the total GFP area and multiplying the ratio by 100. Similarly, LYVE1% is obtained by dividing the total LYVE1 area by the total GFP area and multiplying the ratio by 100.

### Whole-mount immunostaining procedure

All procedures were carried out at 4^°^C on an orbital shaker (Belly Dancer, IBI Scientific) unless otherwise specified. Tissues were harvested and fixed in 2% paraformaldehyde (PFA) overnight. Fixed tissues were washed with Phosphate buffered saline (PBS) thrice for 10 mins each time. The tissues were then permeabilized with PBS + 0.3% Triton X-100 (PBST) for 1 hour after which there blocked with 3% donkey serum in PBST for 2 hours. Primary Antibodies were prepared in PBST, added to the tissue, and allowed to incubate overnight. The next day, the primary antibody was washed off with PBST (5 times of 7 mins each). The secondary and conjugated antibodies were prepared in PBST, added to the tissues, and left to incubate in a black out box in room temperature (RT) for 1.5 hours on an orbital shaker (Belly Button, IBI Scientific). Tissues were then washed 4 times of 10 mins each with PBST. For nuclear visualization, DAPI (Invitrogen) mixed in PBS and added to the tissues and allowed to incubate for 10 mins after which it was washed off with PBS once. The tissue was mounted onto glass slides (Superfrost Plus Microscope slides, Fisherbrand) and covered in ProLong Diamond Antifade Mountant (Invitrogen) and stored at 4°C overnight. The slides were imaged using a Leica SP8 confocal microscope and acquired using Fiji and figures were created using Adobe Photoshop. Primary and secondary antibodies used are listed in supplement table 1.

### Cell culture, *sh*RNA and OSS

Primary hdLECs (PromoCell, C-12216) were cultured in EBM-V2 (PromoCell, C-22121) media on fibronectin (20 μl) coated 6 well plates. Culture passage 6 or lower were used for the knockdown experiments. For the experiment, hdLECs were infected with lentiviral particles either harboring a sh*RNA* against *RICTOR* or a scramble sh*RNA* construct expressing GFP for 48 hours. The hdLECs were then taken off infection and exposed to flow conditions on a test tube rocker (Thermolyne Speci-Mix aliquot mixermodel M71015, Barnstead International) in a 37°C incubator with 5% CO_2_. The lentiviral sh*RICTOR* and shScramble were purchased from VectorBuilder.

### RNA isolation and qRT-PCR

Total RNA was isolated from hdLECs using the RNeasy Plus Mini Kit (Qiagen) following the manufacturer’s instructions. cDNA from the total RNA was synthesized using the Advantage® RT-for-PCR kit (Takara) according to the manufacturer’s protocol. qPCR was performed with Taqman® probes (Applied Biosystems, Thermofisher) in a QuantStudio 3 realtime system (Applied Biosystems). The Cq value of each gene was normalized to the Cq value of GAPDH.

### Western blot

Western bot was performed according to standard protocol. Protein lysates were harvested after OSS treatment for 30 mins or 48h using RIPA buffer (PierceTM, Thermofisher). Protein gel electrophoresis was performed using the Invitrogen mini gel tank, the protein transfer was carried out using the iBlot 2 Dry Blotting system and the antibodies were blotted using the iBind Western Systems. Protein bands were visualized using the SuperSignalTM West Pico PLUS Chemiluminescent Substrate (ThermoFisher).

### Statistics

All data are depicted as mean ± SEM. Data sets comprising of two groups were analyzed using an unpaired 2-sided students *t* test and data sets and significance was achieved at P < 0.05. Data sets containing one independent variable were analyzed using one way analysis of variance (ANOVA) with Tukey’s multiple comparison test to determine significant differences at P < 0.05. GraphPad Prism software (version 9) used to plot quantified data and perform statistical analysis.

### Study Approval

All animal experiments were performed in accordance with the University of South Florida guidelines and were approved by the respective Institutional Animal Care and Use Committee (IACUC) committees. All procedures complied with the standards stated in the “Guide for the Care and Use of Laboratory Animals” (National Institutes of Health, revised 2011).

## Author contributions

RB and YY performed experiments, analyzed data, and wrote the manuscript. LAK, DI, SEB, and JPS performed experiments and edited the manuscript. All authors contributed to and approved the final version of the manuscript.

## Acknowledgments

We thank Dr. Taija Makinen and Dr. Young Kwon Hong for sharing the *Prox1CreER^T2^* mice and the *Prox1-GFP* mice. This work was supported by National Heart, Lung, and Blood Institute (NHLBI) grants R01 HL145397 to Y.Y, R01 HL142905 and R01 HL164825 to J.P.S, AHA Predoctoral Fellowship 827540 to D.I.

## References

1. Breslin, J.W., et al., Lymphatic Vessel Network Structure and Physiology. Compr Physiol, 2018. 9(1): p. 207–299.

2. Yang, Y. and G. Oliver, Development of the mammalian lymphatic vasculature. J Clin Invest, 2014. 124(3): p. 888–97.

3. Koltowska, K., et al., Getting out and about: the emergence and morphogenesis of the vertebrate lymphatic vasculature. Development, 2013. 140(9): p. 1857–70.

4. Petrova, T.V., et al., Defective valves and abnormal mural cell recruitment underlie lymphatic vascular failure in lymphedema distichiasis. Nat Med, 2004. 10(9): p. 974–81.

5. Grada, A.A. and T.J. Phillips, Lymphedema: Pathophysiology and clinical manifestations. J Am Acad Dermatol, 2017. 77(6): p. 1009–1020.

6. Neligan, P.C., T.A. Kung, and J.H. Maki, MR lymphangiography in the treatment of lymphedema. J Surg Oncol, 2017. 115(1): p. 18–22.

7. Kriederman, B.M., et al., FOXC2 haploinsufficient mice are a model for human autosomal dominant lymphedema-distichiasis syndrome. Hum Mol Genet, 2003. 12(10): p. 1179–85.

8. Bazigou, E., et al., Integrin-alpha9 is required for fibronectin matrix assembly during lymphatic valve morphogenesis. Dev Cell, 2009. 17(2): p. 175–86.

9. Ostergaard, P., et al., Mutations in GATA2 cause primary lymphedema associated with a predisposition to acute myeloid leukemia (Emberger syndrome). Nat Genet, 2011. 43(10): p. 929–31.

10. Sabine, A., et al., FOXC2 and fluid shear stress stabilize postnatal lymphatic vasculature. J Clin Invest, 2015. 125(10): p. 3861–77.

11. Ma, G.C., et al., A recurrent ITGA9 missense mutation in human fetuses with severe chylothorax: possible correlation with poor response to fetal therapy. Prenat Diagn, 2008. 28(11): p. 1057–63.

12. Kazenwadel, J., et al., GATA2 is required for lymphatic vessel valve development and maintenance. J Clin Invest, 2015. 125(8): p. 2979–94.

13. Ferrell, R.E., et al., GJC2 missense mutations cause human lymphedema. Am J Hum Genet, 2010. 86(6): p. 943–8.

14. Ostergaard, P., et al., Rapid identification of mutations in GJC2 in primary lymphoedema using whole exome sequencing combined with linkage analysis with delineation of the phenotype. J Med Genet, 2011. 48(4): p. 251–5.

15. Munger, S.J., M.J. Davis, and A.M. Simon, Defective lymphatic valve development and chylothorax in mice with a lymphatic-specific deletion of Connexin43. Dev Biol, 2017. 421(2): p. 204–218.

16. Geng, X., Y.C. Ho, and R.S. Srinivasan, Biochemical and mechanical signals in the lymphatic vasculature. Cell Mol Life Sci, 2021. 78(16): p. 5903–5923.

17. Yang, Y., et al., VE-Cadherin Is Required for Lymphatic Valve Formation and Maintenance. Cell Rep, 2019. 28(9): p. 2397–2412 e4.

18. Cha, B., et al., Mechanotransduction activates canonical Wnt/beta-catenin signaling to promote lymphatic vascular patterning and the development of lymphatic and lymphovenous valves. Genes Dev, 2016. 30(12): p. 1454–69.

19. Zhou, F., et al., Akt/Protein kinase B is required for lymphatic network formation, remodeling, and valve development. Am J Pathol, 2010. 177(4): p. 2124–33.

20. Jhanwar-Uniyal, M., et al., Diverse signaling mechanisms of mTOR complexes: mTORC1 and mTORC2 in forming a formidable relationship. Adv Biol Regul, 2019. 72: p. 51–62.

21. Guertin, D.A., et al., Ablation in mice of the mTORC components raptor, rictor, or mLST8 reveals that mTORC2 is required for signaling to Akt-FOXO and PKCalpha, but not S6K1. Dev Cell, 2006. 11(6): p. 859–71.

22. Bazigou, E., et al., Genes regulating lymphangiogenesis control venous valve formation and maintenance in mice. J Clin Invest, 2011. 121(8): p. 2984–92.

23. Choi, I., et al., Visualization of lymphatic vessels by Prox1-promoter directed GFP reporter in a bacterial artificial chromosome-based transgenic mouse. Blood, 2011. 117(1): p. 362–5.

24. Huang, X.Z., et al., Fatal bilateral chylothorax in mice lacking the integrin alpha9beta1. Mol Cell Biol, 2000. 20(14): p. 5208–15.

25. Altiok, E., et al., Integrin Alpha-9 Mediates Lymphatic Valve Formation in Corneal Lymphangiogenesis. Invest Ophthalmol Vis Sci, 2015. 56(11): p. 6313–9.

26. Saxton, R.A. and D.M. Sabatini, mTOR Signaling in Growth, Metabolism, and Disease. Cell, 2017. 169(2): p. 361–371.

27. Oliver, G., et al., The Lymphatic Vasculature in the 21(st) Century: Novel Functional Roles in Homeostasis and Disease. Cell, 2020. 182(2): p. 270–296.

28. Wang, Y., et al., Smooth muscle cell recruitment to lymphatic vessels requires PDGFB and impacts vessel size but not identity. Development, 2017. 144(19): p. 3590–3601.

29. Kemp, S.S., et al., Defining Endothelial Cell-Derived Factors That Promote Pericyte Recruitment and Capillary Network Assembly. Arterioscler Thromb Vasc Biol, 2020. 40(11): p. 2632–2648.

30. Liu, Y., et al., Edg-1, the G protein-coupled receptor for sphingosine-1-phosphate, is essential for vascular maturation. J Clin Invest, 2000. 106(8): p. 951–61.

31. Yang, X., et al., Angiogenesis defects and mesenchymal apoptosis in mice lacking SMAD5. Development, 1999. 126(8): p. 1571–80.

32. Scallan, J.P., et al., Foxo1 deletion promotes the growth of new lymphatic valves. J Clin Invest, 2021. 131(14).

33. Kume, T., Lymphatic vessel development: fluid flow and valve-forming cells. J Clin Invest, 2015. 125(8): p. 2924–6.

34. Zhang, F., et al., Lymphatic Endothelial Cell Junctions: Molecular Regulation in Physiology and Diseases. Front Physiol, 2020. 11: p. 509.

35. Lu, H. and H. Huang, FOXO1: a potential target for human diseases. Curr Drug Targets, 2011. 12(9): p. 1235–44.

36. Niimi, K., et al., FOXO1 represses lymphatic valve formation and maintenance via PRDM1. Cell Rep, 2021. 37(9): p. 110048.

37. Oh, W.J. and E. Jacinto, mTOR complex 2 signaling and functions. Cell Cycle, 2011. 10(14): p. 2305–16.

38. Sweet, D.T., et al., Lymph flow regulates collecting lymphatic vessel maturation in vivo. J Clin Invest, 2015. 125(8): p. 2995–3007.

39. Sen, B., et al., mTORC2 regulates mechanically induced cytoskeletal reorganization and lineage selection in marrow-derived mesenchymal stem cells. J Bone Miner Res, 2014. 29(1): p. 78–89.

40. Uutela, M., et al., PDGF-D induces macrophage recruitment, increased interstitial pressure, and blood vessel maturation during angiogenesis. Blood, 2004. 104(10): p. 3198–204.

41. Guo, X. and S.Y. Chen, Transforming growth factor-beta and smooth muscle differentiation. World J Biol Chem, 2012. 3(3): p. 41–52.

42. Iivanainen, E., et al., Angiopoietin-regulated recruitment of vascular smooth muscle cells by endothelial-derived heparin binding EGF-like growth factor. FASEB J, 2003. 17(12): p. 1609–21.

43. Wamhoff, B.R., et al., Sphingosine-1-phosphate receptor subtypes differentially regulate smooth muscle cell phenotype. Arterioscler Thromb Vasc Biol, 2008. 28(8): p. 1454–61.

44. Lee, K.S., et al., HB-EGF induces delayed STAT3 activation via NF-kappaB mediated IL-6 secretion in vascular smooth muscle cell. Biochim Biophys Acta, 2007. 1773(11): p. 1637–44.

45. Shiojima, I. and K. Walsh, Role of Akt signaling in vascular homeostasis and angiogenesis. Circ Res, 2002. 90(12): p. 1243–50.

46. Lauffenburger, D.A. and A.F. Horwitz, Cell migration: a physically integrated molecular process. Cell, 1996. 84(3): p. 359–69.

47. Fujio, Y. and K. Walsh, Akt mediates cytoprotection of endothelial cells by vascular endothelial growth factor in an anchorage-dependent manner. J Biol Chem, 1999. 274(23): p. 16349–54.

48. Morales-Ruiz, M., et al., Vascular endothelial growth factor-stimulated actin reorganization and migration of endothelial cells is regulated via the serine/threonine kinase Akt. Circ Res, 2000. 86(8): p. 892–6.

49. Kerr, B.A., et al., Stability and function of adult vasculature is sustained by Akt/Jagged1 signalling axis in endothelium. Nat Commun, 2016. 7: p. 10960.

50. Tonks, N.K., Protein tyrosine phosphatases: from genes, to function, to disease. Nat Rev Mol Cell Biol, 2006. 7(11): p. 833–46.

51. Gelb, B.D. and M. Tartaglia, Noonan syndrome and related disorders: dysregulated RAS-mitogen activated protein kinase signal transduction. Hum Mol Genet, 2006. 15 **Spec No 2**: p. R220-6.

52. Hasegawa, K., et al., Late-onset Lymphedema and Protein-losing Enteropathy with Noonan Syndrome. Clin Pediatr Endocrinol, 2009. 18(3): p. 87–93.

53. Pieper, C.C., et al., MR lymphangiography of lymphatic abnormalities in children and adults with Noonan syndrome. Sci Rep, 2022. 12(1): p. 11164.

54. Ukropec, J.A., et al., SHP2 association with VE-cadherin complexes in human endothelial cells is regulated by thrombin. J Biol Chem, 2000. 275(8): p. 5983–6.

55. Au, A.C., et al., Protein tyrosine phosphatase PTPN14 is a regulator of lymphatic function and choanal development in humans. Am J Hum Genet, 2010. 87(3): p. 436–44.

56. Aboujaoude, W., M.L. Milgrom, and M.V. Govani, Lymphedema associated with sirolimus in renal transplant recipients. Transplantation, 2004. 77(7): p. 1094–6.

57. Romagnoli, J., et al., Severe limb lymphedema in sirolimus-treated patients. Transplant Proc, 2005. 37(2): p. 834–6.

58. Ersoy, A. and N. Koca, Everolimus-induced lymphedema in a renal transplant recipient: a case report. Exp Clin Transplant, 2012. 10(3): p. 296–8.

59. Paik, J.H., et al., FoxOs are lineage-restricted redundant tumor suppressors and regulate endothelial cell homeostasis. Cell, 2007. 128(2): p. 309–23.

60. Sabine, A., et al., Mechanotransduction, PROX1, and FOXC2 cooperate to control connexin37 and calcineurin during lymphatic-valve formation. Dev Cell, 2012. 22(2): p. 430–45.

61. Tatin, F., et al., Planar Cell Polarity Protein Celsr1 Regulates Endothelial Adherens Junctions and Directed Cell Rearrangements during Valve Morphogenesis. Dev Cell, 2013. 26(1): p. 31–44.

62. Rouhani, S.J., et al., Roles of lymphatic endothelial cells expressing peripheral tissue antigens in CD4 T-cell tolerance induction. Nat Commun, 2015. 6: p. 6771.

